# A Hybrid Deep Clustering Approach for Robust Cell Type Profiling Using Single-cell RNA-seq Data

**DOI:** 10.1101/511626

**Authors:** Suhas Srinivasan, Nathan T. Johnson, Dmitry Korkin

## Abstract

Single-cell RNA sequencing (scRNA-seq) is a recent technology that enables fine-grained discovery of cellular subtypes and specific cell states. It routinely uses machine learning methods, such as feature learning, clustering, and classification, to assist in uncovering novel information from scRNA-seq data. However, current methods are not well suited to deal with the substantial amounts of noise that is created by the experiments or the variation that occurs due to differences in the cells of the same type. Here, we develop a new hybrid approach, Deep Unsupervised Single-cell Clustering (DUSC), that integrates feature generation based on a deep learning architecture with a model-based clustering algorithm, to find a compact and informative representation of the single-cell transcriptomic data generating robust clusters. We also include a technique to estimate an efficient number of latent features in the deep learning model. Our method outperforms both classical and state-of-the-art feature learning and clustering methods, approaching the accuracy of supervised learning. The method is freely available to the community and will hopefully facilitate our understanding of the cellular atlas of living organisms as well as provide the means to improve patient diagnostics and treatment.

## Introduction

Despite the centuries of research, our knowledge of the cellular architecture of human tissues and organs is still very limited. Microscopy has been conventionally used as a fundamental method to discover novel cell types, study cell function and cell differentiation states through staining and image analysis [1]. However, this approach is not able to identify heterogeneous sub-populations of cells, which might look similar, but perform different functions. Recent developments in single-cell RNA sequencing (scRNA-seq) have enabled harvesting the gene expression data from a wide range of tissue types, cell types, and cell development stages, allowing for a fine-grained discovery of cellular subtypes and specific cell states [2]. Single-cell RNA sequencing data have played a critical role in the recent discoveries of new cell types in the human brain [3], gut [4], lungs [5], and immune system [6], as well as in determining cellular heterogeneity in cancerous tumors, which could help improve prognosis and therapy [7, 8].

Single-cell experiments produce datasets that have three main characteristics of big data: volume (number of samples and number of transcripts per each sample), variety (types of tissues and cells), and veracity (missing data, noise, and dropout events) [9]. Recently emerging large initiatives, such as the Human Cell Atlas [10], rely on single-cell sequencing technologies at an unprecedented scale, and have generated datasets obtained from hundreds of thousands and even millions of cells. The high numbers of cells, in turn, allow to account for data variability due to cellular heterogeneity and different cell-cycle stages. As a result, there is a critical need to automate the processing and analysis of scRNA-seq data. For instance, for the analysis of large transcriptomics datasets, computational methods are frequently employed that find patterns associated with the cellular heterogeneity or cellular development, and group cells according to these patterns.

If one assumes that all cellular types or stages extractable from a single-cell transcriptomics experiment have been previously identified, it is possible to apply a supervised learning classifier. The supervised learning methods are trained on the data extracted from the individual cells whose types are known. The previously developed approaches for supervised cell type classification have leveraged data from image-based screens [11] and flow cytometry experiments [12]. There has also been a recent development of supervised classifiers for single-cell transcriptomic data [13]. While a supervised learning approach is expected to be more accurate in identifying the previously observed cellular types, its main disadvantage is the limited capacity in discovering new cell types or identifying the previously known cell types whose RNA-seq profiles differ from the ones observed in the training set.

Another popular technique for scRNA-seq data analysis is unsupervised learning, or clustering. In this approach, no training data are provided. Instead, the algorithm looks to uncover the intrinsic similarities shared between the cells of the same type and not shared between the cells of different types [14]. Often, the clustering analysis is coupled with a feature learning method to filter out thousands of unimportant features extracted from the scRNA-seq data. In a recent study, the Principal Component Analysis (PCA) approach was used on gene expression data from scRNA-seq experiments profiling neuronal cells [15]. With the goal of identifying useful gene markers that underlie specific cell types in the dorsal root ganglion of mice, 11 distinct cellular clusters were discovered. Other approaches have also adopted this strategy of combining a simple, but efficient feature learning method with a clustering algorithm, to detect groups of cells that could be of different sub-types or at different stages in cellular development [21, 65]. One challenge faced by such an approach is due to scRNA-seq data exhibiting complex high-dimensional structure, and such complexity cannot be accurately captured by fewer dimensions when using simple linear feature learning methods.

A nonlinear method frequently used in scRNA-seq data analysis for clustering and visualization is t-distributed stochastic neighbor embedding (t-SNE) [16]. While, t-SNE can preserve the local clusters, preserving the global hierarchical structure of clusters is often problematic [17]. Furthermore, the conventional feature learning methods may not be well suited for scRNA-seq experiments that have considerable amount of both experimental and biological noise or the occurrence of dropout events [18, 19]. To address this problem, two recent methods have been introduced, pcaReduce [20] and SIMLR [21]. pcaReduce integrates an agglomerative hierarchical clustering with PCA to generate a hierarchy where the cluster similarity is measured in subspaces of gradually decreasing dimensionalities. The other approach, SIMLR, learns different cell-to-cell distances through by analyzing the gene expression matrix; it then performs feature learning, clustering, and visualization. The computational complexity of the denoising technique in SIMLR prevents its application on the large datasets. Therefore, a different pipeline is used to handle large data, where the computed similarity measure is approximated, while the diffusion approach to reduce the effects of noise is not used. In addition to the dimension reduction methods, K-means is a popular clustering method used in single-cell transcriptomics analysis. While being arguably the most popular divisive clustering algorithm it has several limitations [22, 23].

In this work, we looked at the possibility to leverage an unsupervised deep learning approach [24] to handle the complexities of scRNA-seq data and overcome the limitations of the current feature learning methods. It has been theoretically shown that the multilayer feed-forward artificial neural networks, with an arbitrary squashing function and sufficient number of hidden units (latent features) are the universal approximators [25] capable of performing the dimensionality reduction [26]. Here, we propose the use of denoising autoencoder (DAE) [27], an unsupervised deep learning architecture that has previously proven successful for several image classification [28] and speech recognition [29] tasks. DAEs are different from other deep learning architectures in their ability to handle noisy data and construct robust features. We add a novel extension to the DAE called **D**enoising **A**utoencoder **W**ith **N**euronal approximator (DAWN), which decides the number of latent features that is required to represent efficiently any given dataset. To overcome the limitations of K-means clustering, we integrate our DAWN approach with the expectation-maximization (EM) clustering algorithm [30]. We use the features generated by DAWN as an input to the EM clustering algorithm and show that our hybrid approach has higher accuracy when compared to the traditional feature learning and clustering algorithms discussed above. In particular, we are able to recover clusters from the original study without using any knowledge about the tissue-specific or cell type specific markers. As a result, our hybrid approach, **D**eep **U**nsupervised **S**ingle-cell **C**lustering (DUSC), helps to overcome the noise in the data, captures features that are representative of the true patterns, and improves the clustering accuracy.

## Methods

### Overview of the approach

The goal of this work is to design a method capable of identifying cellular types from single-cell transcriptomics data of a heterogenous population without knowing *a priori* the number of cell types, sub-population sizes, or the gene markers of the population. Our hybrid approach, DUSC, combines deep feature learning and expectation-maximization clustering. The feature learning leverages the denoising autoencoder (DAE) and includes a new technique to estimate the number of required latent features. To assess the accuracy of our approach, we test it on a series of scRNA-seq datasets that are increasingly complex with respect to the biological and technical variability. The performance of our method is then compared with performances of classical and state-of-the-art unsupervised and supervised learning methods.

The DUSC computational pipeline consists of four main stages (Fig. 1). Following a basic data quality check, we first pre-process the data for training DAE. Second, we perform feature learning using DAWN, which includes training DAE and hyper-parameter optimization. We note that the data labels are not required during the training part of the pipeline; instead, the labels are used solely to test the accuracy of DUSC across the datasets and to compare it against the other methods. Third, we use the previously published four feature learning methods, Principal Component Analysis (PCA) [31], Independent Component Analysis (ICA) [32], t-SNE [16], and SIMLR [21], to generate the compressed dimensions for the same scRNA-seq dataset that was used as an input to DAWN. This allows us to assess how well the autoencoder learns the latent features compared to the other methods. Finally, we use the reduced feature representations from each of the above five methods and pass them as an input to the two clustering algorithms, K-means (KM) and expectation maximization (EM), to assess the clustering accuracy.

**Figure 1.**
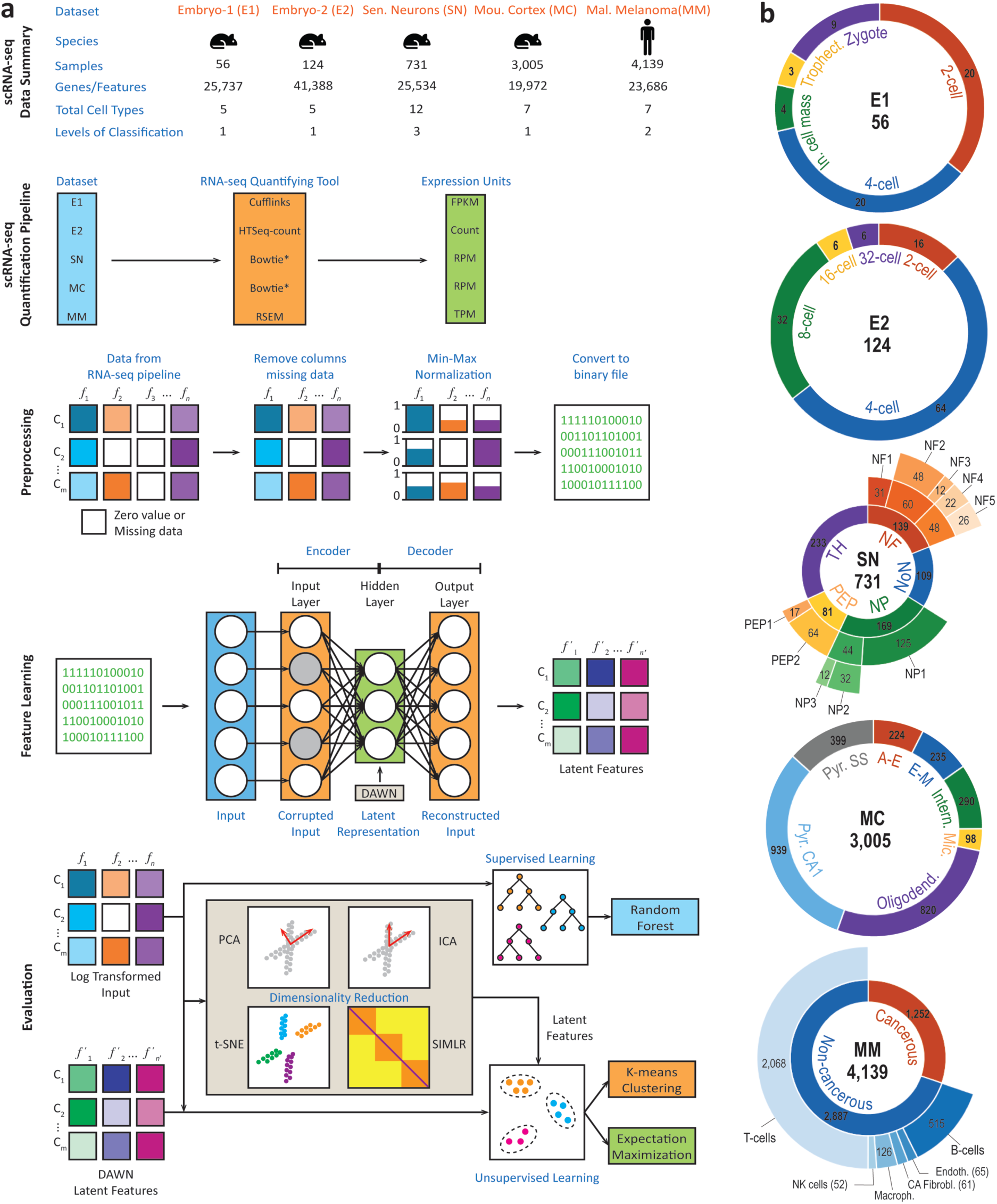
Overview of DUSC approach. (a) Basic stages of the deep clustering method and overview of the five datasets it was applied to. Each of the five datasets was processed using a different RNA-seq quantification tool, with the data quantified in different expression units. During the evaluation, our approach was compared against the standard clustering methods as well as their enhanced versions using four feature learning approaches. (b) The detailed description of the five datasets: Embryonic Dataset-1 (E1), Embryonic Dataset-2 (E2), Sensory Neurons (SN), Mouse Cortex (MC), and Malignant Melanoma (MM), their multi-level hierarchical organizations, and subpopulation distribution. The total number of cell samples is depicted in the center of each sunburst chart.

### Denoising autoencoder model design

During the data pre-processing stage, a dataset is defined as a matrix where the rows correspond to the cell samples, and the columns correspond to the feature vectors containing the gene expression values. To reduce the computational complexity, we remove the matrix columns where all values are zeros, which is the only type of gene filtering used in this method (number of genes removed in each dataset are detailed in Sup. Table S1). This minimal filtering procedure is different from a typical gene filtering protocol, whose goal is to restrict the gene list to a few hundred or a few thousand genes [15, 49]. Here, we aim to provide as much data as possible for our deep learning algorithm to capture the true data structure. The columns are then normalized by scaling the gene expression values to [0,1] interval:

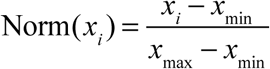

where *x*_max_ and *x*_min_ are the maximum and minimum values across all feature values in vector *x* respectively, and *x*_*i*_ is a feature value in *x.* The normalized matrix is converted from a 64-bit floating point representation to a 32-bit representation for native GPU computation and then to a binary file format, to reduce the input-output costs and GPU memory usage during the computation.

An autoencoder [24] is a type of artificial neural network that is used in unsupervised learning to automatically learn features from the unlabeled data. A standard neural network is typically designed for a supervised learning task and includes several layers where each layer consists of an array of basic computational units called neurons, and the output of one neuron serves as an input to another. The first, input, layer takes as an input a multi-dimensional vector *x*^(*i*)^ representing an unlabeled example. The intermediate, hidden, layers are designed to propagate the signal from the input layer. The last, output, layer calculates the final vector of values *z*^(*i*)^ corresponding to the class labels in a supervised learning setting. In the autoencoder, the output values are set to be equal to the input values, *x*^(*i*)^ = *z*^(*i*)^, and the algorithm is divided into two parts, the encoder and the decoder. In the encoder part, the algorithm maps the input to the hidden layer’s latent representation *y* = *s*(*Wx* + *b*), where *s*(*x*) is a sigmoid function: 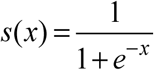. In the decoder part, the latent representation *y* is mapped to the output layer: *z* = *s*’(*W*’*y* + *b*’). As a result, *z* is seen as a prediction of *x,* given *y.* The weight matrix, *W*’, of the reverse mapping is constrained to be the transpose of the forward mapping, which is referred to as tied weights given by *W*’ =*W*^*T*^ The autoencoder is trained to minimize an error metric defined as the cross-entropy of reconstruction, *L*_*H*_ (*x, z*), of the latent features:

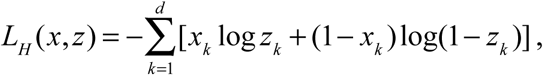

where *d* is the length of the feature vector.

To prevent the hidden layer from simply learning the identity function and forcing it to discover more robust features, a DAE is introduced. A DAE is trained to recover the original input from its corrupted version [27]. The corrupted version is obtained by randomly selecting *n*_*d*_ features of each input vector *x*^(*i*)^ and assigning them zero values. This stochastic process is set up by 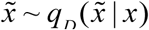, where 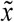 is the corrupted input. Even when the corrupted vectors are provided to the neural network, the reconstruction error is still computed on the original, uncorrupted, input. The optimal number of hidden layers and hidden units for the DAE in this approach is explored as a part of model optimization. The DAE is implemented using the Theano Python library [33], which supports NVidia CUDA. This implementation allows for fast training of the neural network layers with large numbers of nodes using NVidia GPUs.

### Model optimization

The overall architecture of the DAE implemented in our approach consists of an input layer, an output layer, and one hidden layer. There are multiple parameters in this DAE architecture that can be optimized. The task of hyperparameter optimization is fairly unambiguous for supervised learning problems [63], where the data are labeled, and a neural network can be tuned to set its many parameters such that it achieves an optimal classification performance (*e.g.*, measured by accuracy, f-measure, or other measures). However, in the case of unsupervised clustering where no labeled data are provided and the neural network parameters are optimized to minimize the reconstruction error, the impact of this error metric on clustering is not known *a priori*. To make this optimization a computationally feasible task, we focus on tuning the number of hidden units, which is expected to have the most significant impact on the model performance [64], given its single hidden-layer architecture. The tuning is performed by adopting the ideas from Principal component analysis (PCA) [31].

PCA works by converting the initial set of features, which potentially correlate with each other, into linearly uncorrelated features (principal components), through an orthogonal transformation of the feature space. It has been shown that PCA is a special case of the autoencoder where a single hidden layer is used, the transformation function in the hidden units is linear, and a squared error loss is used [28]. PCA offers an automated technique to select the first *n* principal components required to capture a specified amount of variance in a dataset [34], *i.e.* in a linear autoencoder the principal components are simply the nodes in the hidden layer. The similarity between the two approaches leads us to test, if one can use PCA to approximate the number of hidden units required in an autoencoder to capture most of the data complexity in each dataset. As a result, we apply PCA immediately preceding DAE, using the original dataset as an input to PCA and producing, as an output, the number *n* of principal components required to capture 95% of the dataset variance (PCA for all datasets is shown in Sup. Fig. S2). The same data are then processed by DAE with the number of hidden units set to *n.* We then assess if this additional optimization stage to DAE improves the performance of our approach and call this new extension as **D**enoising **A**utoencoder **W**ith **N**euronal approximation (DAWN).

Since we focus only on the impact of the number of hidden units on the learning efficiency, the settings for all other parameters of the DAE are selected based on a recent work that used DAEs to learn important features in a breast cancer gene expression dataset [35]. Specifically, we set: (1) the learning rate to 0.05; (2) training time to 500 epochs, which has been reported to be sufficient for the reconstruction error to converge; (3) batch size to 20, to limit the number of batches for the larger datasets; and (4) corruption level to 0.1, which specifies that 10% of the input vector features are randomly set to zeroes. The number of hidden neurons estimated for each dataset are provided in Sup. Table S7.

To generate cell clusters from the learnt features of DAWN, we use the Expectation-Maximization (EM) clustering algorithm [30]. We choose this clustering method because it overcomes some of the main limitations of K-means, such as sensitivity to initial clustering, instance order, noise and the presence of outliers [23]. In addition, EM is a statistical-based clustering algorithm that can work with the clusters of arbitrary shapes and is expected to provide clustering results that are different to those ones of K-means, which is a distance-based algorithm and works best on the compact clusters. Finally, EM clustering can estimate the number of clusters in the dataset, while K-means requires the number of clusters to be specified as an input. These attributes make EM algorithm a good candidate, because we expect the latent features of DAWN to have specific distributions corresponding to different groups of cells, and we can also approximate the number of clusters.

### Comparative assessment of DAWN against existing feature learning approaches

The assessment of the overall performance of our DUSC pipeline includes evaluating the performances of both, the DAWN method and EM clustering algorithm. To evaluate the accuracy of feature learning by DAWN, we compare it against the four other feature learning methods: a stand-alone PCA, ICA, t-SNE, and SIMLR.

PCA is widely used across many areas including scRNA-seq analysis to reduce the data complexity and to make the downstream analysis computationally more feasible. During the assessment stage, we set PCA algorithm to select the minimal number of principal components required to learn 95% of variance in the data. Independent component analysis (ICA) is another statistical method designed to separate a multivariate signal into additive subcomponents, which has been applied to a wide range of image analysis and signal processing tasks [36, 37]. Assuming that the scRNA-seq data can be represented as following a mixture of non-Gaussian distributions, ICA can potentially determine the individual independent components that best capture the cell type information in the transcriptomics data. ICA has been previously successfully applied to scRNA-seq data where changes in the transcriptome were resolved to an improved temporal resolution during cell differentiation [38]. However, both PCA and ICA make certain assumptions on the data structure: in addition to being linear methods, they are not designed to handle the considerable amount of noise present in the scRNA-seq data. Unlike PCA, the ICA algorithm cannot automatically choose the number of components required to learn a given amount of data variance. Hence, we manually set the number of components to the same number derived by the PCA method when it is required to learn 95% of the data variance.

t-Distributed stochastic neighbor embedding (t-SNE) is a nonlinear feature learning technique specifically designed to map and visualize high-dimensional data into two-dimensional (2D) or three-dimensional (3D) spaces [16]. t-SNE is often used in scRNA-seq studies to visualize cell subpopulations in a heterogeneous population [39]. The technique is very efficient in capturing critical parts of the local structure of the high-dimensional data, while facing difficulties in preserving the global hierarchical structure of clusters [17]. Another potential drawback of t-SNE is the time and space complexities that are both quadratic in the number of samples. Thus, this method is typically applied to a smaller subset of highly variable gene features. When evaluating it against DAWN, we use t-SNE only for the feature learning. t-SNE is dependent on an important parameter, perplexity, which estimates the effective number of neighbors for each data point. Here, instead of setting it arbitrarily in the range of [5,50], we calculate it precisely for each dataset based on Shannon entropy (discussed below).

Single-cell interpretation via multi-kernel learning (SIMLR) is a recent state-of-the-art computational approach that performs the feature learning, clustering, and visualization of scRNA-seq data by learning a distance metric that best estimates the structure of the data [21]. The general form of the distance between cells *i* and *j* is expressed as a weighted combination of multiple kernels:

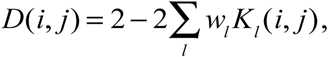

where, *w*_*l*_ is the linear weight value, which represents the importance of each individual kernel *K*_*l*_ (*i, j*), and each kernel is a function of the expression values for cells *i* and *j.* The similarity matrix *S*_*ij*_ is therefore a *N* × *N* matrix where *N* is the number of samples, capturing the pairwise expression-based similarities of cells:

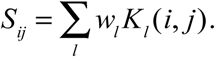

In SIMLR, to reduce the effects of noise and dropouts in the data, a diffusion-based technique [40] is employed. However, this technique is computationally expensive and therefore can be only applied to small or medium size datasets (*e.g*., in a previous work, any dataset with a sample size greater than 3,000 did not use this technique [21]). Hence, the noise and dropouts effects remain present in the large datasets. Furthermore, the SIMLR framework uses K-means as its clustering algorithm and is affected by the previously discussed limitations. While SIMLR has the capability to estimate the number of clusters, to compare DAWN with the best possible performance of SIMLR, we set the true number of clusters for each dataset as an input to SIMLR. Note that this information about the number of clusters is not provided to any other method. The PCA, ICA, and t-SNE algorithms were evaluated using the implementations in the Python scikit-learn library [41], while SIMLR was evaluated using its implementation as an R package.

### Evaluation protocol

All five feature learning methods are evaluated by integrating each of them with one of the two clustering algorithms used in this work, K-means or EM. To do so, we use the latent features uncovered by each of the five methods as inputs to the two clustering algorithms. This setup also allows to comparatively assess the individual contributions towards the prediction accuracy by each of DUSC’s two components, DAWN and EM clustering. Indeed, one can first assess how much the addition of DAWN to K-means or EM can affect the clustering accuracy by comparing the performance with K-means and EM when using the default features. Second, one can determine if the EM-based hybrid clustering approach is more accurate than K-means based approach for each of the five feature learning methods (including DAWN). In total, we evaluate all 5×2=10 combinations of hybrid clustering approaches.

Alternatively, to determine if the neuronal approximation implemented in DAWN improves a standard DAE, we compared the performance of DUSC pipeline with DAWN and with two DAE configurations. Although, the number of hidden units of a DAE can be set to any arbitrary value, we manually set it to 50 in the first configuration and 100 in the second one, making these configurations computationally feasible [35] (which we name as DAE-50 and DAE-100 for convenience).

Finding the optimal number of clusters in a dataset is often considered an independent computational problem. Therefore, for the assessment of clustering accuracy, we set the expected number of clusters to be the number of cell types originally discovered in each study. To establish the baseline, we applied KM and EM clustering on the original datasets with zero-value features filtered out, and the data being log_10_ transformed. The KM and EM methods are implemented using WEKA package [42].

After evaluating the performance of DUSC against other unsupervised methods, we next compare it against a state-of-the-art supervised learning approach. While a supervised learning method is unable to discover new cell types, it is expected to be more accurate in identifying the previously learned types that the algorithm has been trained on. We use the log-transformed data as an input and apply the multi-class Random Forest (RF) algorithm [43] implemented in WEKA, with a 10-fold cross validation protocol [44] that selects the best model with the highest accuracy.

For each of the above evaluations, it is desirable to have a common evaluation metric that can handle multi-class datasets. Here, we use a simple accuracy measure (*Acc*), which can be calculated by comparing the predicted cell clusters with the known cell labels:

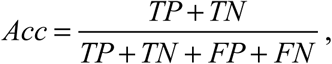

where *TP* is the number of true positives, *TN* is the number of true negatives, *FP* is the number of false positives, and *FN* is the number of false negatives.

In addition to the standard evaluation of the performance accuracy, the following three characteristics of the method’s performance are explored. First, we study performance of the methods as a function of data complexity. Each of the five datasets considered in this work varies with respect to the sample size distributions across different cell types, numbers of cell types, and cell type hierarchy. These three properties are expected to affect the complexity of cluster separation, prompting one to study the correlation between these properties and the clustering accuracy. To measure the distribution balance of samples across all cell types for each dataset we use normalized Shannon Entropy [18]:

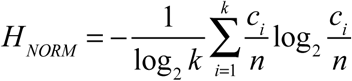

where *n* is the total number of samples, *k* is the number of cell types, and *c*_*i*_ is the number of samples in cell type *i.* Thus, *H*_*NORM*_ approaches 0 if the dataset is unbalanced and 1 if it is balanced.

Second, since the learned latent features are designed to capture the complexity of each dataset and create its reduced representation, one can assess the data compression performance of DAWN. The data compression ratio (*CR*) is defined as a ratio between the sizes of the original uncompressed and compressed datasets. A normalized value that allows interpreting the compression performance more intuitively in terms of feature space compressed (*FSC*), is defined as:

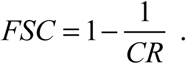

*FSC* value approaching 1 implies that the original dataset has been compressed to a very small feature set size.

Finally, to determine if DUSC can improve the cell type cluster visualization, we generate two-dimensional embeddings by applying t-SNE to the features of the four previously considered feature learning methods as well as features generated by DAWN. In our qualitative assessment of the visualizations, we expect to see the clusters that are well-separated and compact (*i.e.*, the intra-cluster distances are much smaller than inter-cluster distances), and the instances of incorrect clustering are rare.

## Results

### Datasets

For the assessment of our approach, we chose five single-cell RNA-seq datasets (Fig. 1): Embryonic Dataset-1 (E1), Embryonic Dataset-2 (E2), Sensory Neurons (SN), Mouse Cortex (MC), and Malignant Melanoma (MM) [46-50]. These datasets were selected to represent areas where scRNA-seq technology had significant impact [51]. The areas included embryonic development, cellular heterogeneity in the nervous system and cellular heterogeneity in a disease (cancer). The datasets originated from a model organism (mouse) and human. In total, 8,055 single-cell samples were analyzed (Fig. 1a). All datasets were downloaded in a quantified format from publicly available sources listed in the studies [52, 53, 54, 55]. To test the scalability and robustness of the proposed method, the datasets were chosen such that they exhibited variability across multiple parameters: (a) number of sequenced cells (from 56 cells to 4,139 cells), (b) number of genes quantified (from ∼19,000 to ∼41,000 genes), (c) different sequencing and quantification pipelines, (d) cellular heterogeneity during development or disease, and (e) varying cellular hierarchy and number of cellular types (with 1 to 3 levels of hierarchy and 5 to 12 cell types/subtypes). Many of the cellular types include subpopulations corresponding to the cellular subtypes (Fig. 1b). Specifically, the cellular subtypes in SN and MM datasets are hierarchically organized; SN has a three-level hierarchy, while MM has two levels. The distributions of number of genes quantified per cell varied significantly: for E1 and E2, the distribution was centered around ∼13,000 genes, and for SN, MC and MM the distributions were centered around ∼4,000 genes (Sup. Fig. S1). Using the normalized Shannon entropy, *H*_*NORM,*_ we found that the distribution balance of samples across cell types also varied, with the first level of SN being the most balanced and the second level of MM being the most unbalanced sets, correspondingly (see also **Effects of data balance on accuracy** subsection).

### Comparison with clustering and classification algorithms

We first evaluated the overall performance of our clustering approach, DUSC, and its most critical part, a new feature learning method DAWN. To test if DUSC could improve the discovery of cell type clusters in scRNA-seq data, we compared the clustering of our hybrid approach with (i) clustering that had no feature selection, and (ii) the same clustering methods that now employed the classical and state-of-the-art unsupervised feature learning methods. We expected that clustering with no feature selection would perform the worst, thus establishing a baseline for comparative analysis. We also assessed the classification accuracy of Random Forest (RF), a state-of-the-art supervised learning algorithm. The latter approach represents the best-case scenario when all cell types are known.

For E1 dataset, all methods except KM recovered the clusters with similarly high accuracies. As expected, the small sample size and a few cell types to consider made the clustering a simpler task (Fig. 2a and Sup. Table S2). When processing E2, DUSC had the highest accuracy among all methods, and while there was only small accuracy drop for RF, both KM and EM experienced significant losses in accuracy. The drop in performance on E2 dataset, which had the same number of cell types as E1, could be explained by the fact that both the sample size and feature size approximately doubled, therefore quadrupling the problem size and making it a harder computational challenge. For the main hierarchy level in SN (SN-i), the sample size was 731, making it a larger search space, but with only five major cell types. Here, DUSC performed well (*Acc*=0.9) and was closely followed by RF, while KM and EM performed poorly (accuracies were 0.88, 0.53, and 0.62 correspondingly). For the second level of SN subtypes (SN-ii), the sample size was still 731, but the number of cell subtypes increased to 9, thus resulting in a smaller sample size for each cell subtype (Fig. 1b) and smaller feature differences between the subtypes. As a result, it was not surprising that all methods experienced a drop in their performance, with RF performing best (*Acc*=0.74) and DUSC being the first among the unsupervised methods, closely behind RF (*Acc*=0.69). When considering the lowest level of SN, SN-iii, with the number of subtypes being 12 and cell cluster sizes ranging from 12 to 233, we noticed that RF and DUSC both have similar accuracy (*Acc*=0.71), while KM and EM still performed poorly (*Acc*=0.46 and 0.40, correspondingly). We note that for all evaluations based on the subtypes of either SN-i (*i.e.*, SN-ii and SN-iii levels) or MM-i (*i.e.*, MM-ii level), we did not filter out the major cell clusters from the higher levels to recursively process sub-clusters of the lower levels. This is because, the cellular hierarchy was not known *a priori* when analyzing a novel dataset and its structure could only be discovered after the recursive analysis of sub-clusters that, in turn, required multiple iterations. Here, we generated clusters only through a single pass of our processing pipeline.

**Figure 2.**
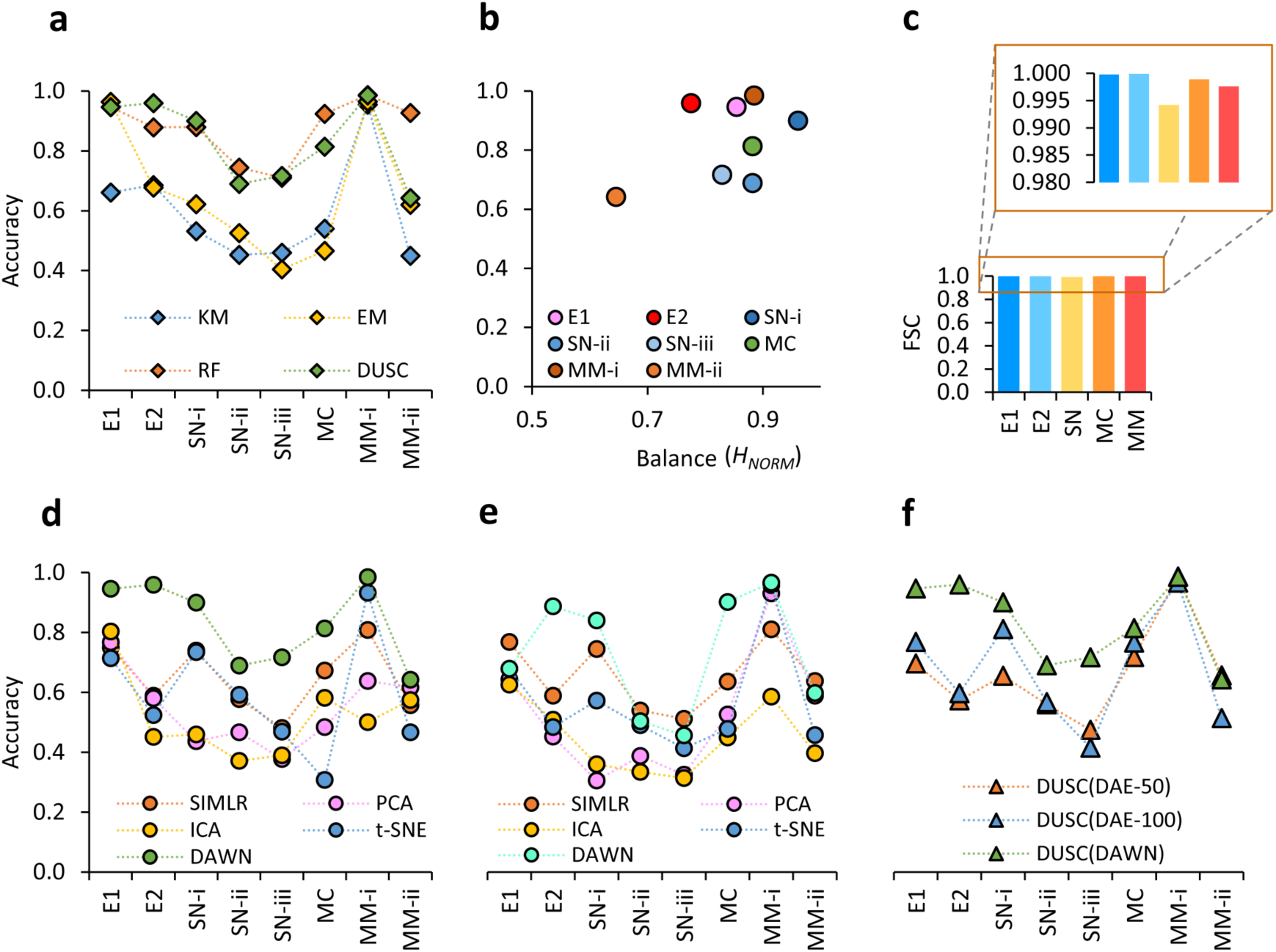
Comparative assessment of DUSC. The methods considered in this figure include: K-means (KM), Expectation-Maximization (EM), Random Forest (RF), Principle Component Analysis (PCA), Independent Component Analysis (ICA), t-Distributed stochastic neighbor embedding (t-SNE), Single-cell Interpretation via Multikernel Learning (SIMLR) and Deep Unsupervised Single-cell Clustering (DUSC). The datasets used in the figure include Embryonic Dataset-1 (E1), Embryonic Dataset-2 (E2), Sensory Neurons (SN-i, SN-ii, and SN-iii correspond to the subpopulations at the first, second, and third levels of hierarchy, respectively), Mouse Cortex (MC), and Malignant Melanoma (MM-i and MM-ii correspond to the subpopulations at the first and second levels of hierarchy, respectively). (a) Overall performance of DUSC in comparison with two clustering approaches, KM and EM, and a state-of-the-art supervised learning approach, RF. DUSC outperforms both clustering methods, and its accuracy is comparable with that of the supervised classifier; (b) The performance accuracy by DUSC is affected by the distribution balance of the subpopulations forming the dataset: applying DUSC to the more unbalanced dataset result in the lower accuracy and vice versa (c) feature space compressed (FSC) calculated for all five datasets; (d) The performance of EM clustering combined with DAWN and other feature learning methods. DAWN shows a significantly greater improvement compared to the other methods; (e) The performance of K-means clustering combined with DAWN and other feature learning methods. Similar to (d), DAWN shows a superior improvement of the stand-alone clustering approach; (f) The clustering performance of DUSC using DAWN, versus using two manual configurations of the standard DAE (DAE-50 and DAE-100). DAWN performs significantly better than the manual configurations and with fewer hidden neurons.

The size of the next dataset, MC, was several folds greater than of the previous two, and in this case, RF had the best accuracy (*Acc*=0.92) and our unsupervised method DUSC had a significantly higher accuracy (*Acc*=0.81) than KM and EM (*Acc*=0.54 and 0.57, correspondingly). Lastly, in the final dataset, MM, we initially tried to find only two clusters of cancerous and non-cancerous cells, and this binary problem with two very different cell types and approximately the same cluster sizes was unsurprisingly an easy challenge. Thus, all methods perform very well with the accuracies above 0.95, but DUSC still lead the unsupervised algorithms with the same accuracy as RF (*Acc*=0.99). When the subtypes of non-cancerous cells had to be considered as separate groups along with cancerous cells (MM-ii), the complexity of the problem increased, and all unsupervised algorithms experienced a significant drop in performance when compared to RF (*Acc*=0.93), with DUSC still achieving the best result (*Acc*=0.64).

In summary, the assessment on all four datasets demonstrated that DUSC performed better than the KM and EM clustering algorithms and in many instances by large margins. Even more importantly, DUSC had a comparable performance with Random Forest supervised approach in many cases, and in some cases even outperformed it.

### Effects of data balance on accuracy

The data balance metric introduced in this work allowed us to find how the data complexity and imbalance affected the performance of DUSC (Fig. 2b). Indeed, for all unsupervised methods, including DUSC, the clustering accuracy was impacted by the data complexity (Sup. Fig. S3 and Sup. Table S3). This was especially evident in the cases of SN-ii and MM-ii datasets, where the number of cell types increased compared to the original datasets, SN-i and MM-i, respectively. The higher number of clusters, in turn, lead to the higher variation in cluster sizes, and in the same time, lower number of differentiating features. Here, we observed that both data balance and clustering accuracy decreased when moving down the cell type hierarchy in SN and MM datasets.

### Feature compression

To study the information content of the initially sparse feature space, another metric, feature space compressed (FSC), was used for DUSC (Fig. 2c). With the combination of the pre-processing stage and the neuronal approximation, DAWN compressed at least 0.994 (99.4%) of the original feature space reaching 0.998 (99.8%) for four out of five datasets (Sup. Table S7). The maximum compression occurred for E2, where 41,388 of the original features were cleaned and compressed to just three latent features resulting in FSC of 99.99%. The data compression capacity of DAWN could also be a useful tool for storing cell-type critical information in large scRNA-seq studies. For instance, the size of an average dataset obtained from a single study could be reduced from 1 gigabyte to only 5 megabytes using DAWN. We note that the highly efficient compression occurred simultaneously when improving the clustering performance.

### Assessment of unsupervised feature learning algorithms and their impact on clustering

We next compared the performance of DUSC against the four feature learning methods, SIMLR, PCA, ICA and t-SNE. Since DUSC is a hybrid approach that combined a new feature learning method (DAWN) and a clustering algorithm (EM), for a fair comparison, we also paired the other four feature learning methods with EM clustering method (Fig. 2d and Sup. Table S4). The results showed that the previously observed effects of the sample size, number of cell clusters, and number of important features on DAWN’s performance also affected the other four methods. For the easier datasets w.r.t the above criteria, such as E1 and MM-i, all the algorithms had the accuracy greater than 0.7, with DUSC reaching significantly higher accuracies of 0.95 and 0.99, respectively. Interestingly, when more complex problems were considered, *i.e.*, E2 and MC, we noticed a significant performance drop for all algorithms; however when compared to SIMLR, the best performing method of the four currently existing ones, DUSC still clustered E2 more accurately (*Acc*=0.96) and also had a 14% higher accuracy (*Acc*=0.81) on MC dataset. We also recall that SIMLR was used in a less challenging setup when the true number of clusters was provided as an input. Overall, DUSC had the better accuracy across all datasets, compared to all other unsupervised feature learning algorithms.

### Assessment of the contributing factors in the hybrid approach

We then hypothesized that between the feature learning (DAWN) and clustering (EM) components of our approach, DAWN was contributing more to the clustering accuracy. To determine the impact of DAWN, we paired it as well as the four other feature learning methods with K-Means clustering. We found that DAWN either exceeded the clustering accuracy of SIMLR, (for E2, SN-i, MC and MM-i) or closely matched it in the other cases (Fig. 2e and Sup. Table S5). The other methods, PCA, ICA and t-SNE had significantly lower accuracies for the majority of the tasks. The findings suggested that DAWN provided the key contribution towards improving the clustering accuracy. A consistent trend that was observed across all methods (Figs. 2d,e) was that for SN and MM datasets, the accuracy decreased as the feature learning and clustering methods traversed the cell type hierarchies. The smaller differences in the numbers of uniquely expressed genes together with a larger set of common genes across the cellular subtypes, compared to the main cellular types, made it a more challenging problem for the feature learning and clustering.

### Assessment of neuronal approximation

To further assess the benefits of our novel neuronal approximation in DAWN, we compared it with the standard DAE. We created two configurations of the standard DAE, by choosing the number for the hidden units to be 50 and 100 respectively. All other aspects of the DUSC approach were kept intact, and the end-to-end analysis was repeated for DAE-50 and DAE-100. The clustering results showed that DAWN outperformed the standard DAE configurations in six cases and had extremely similar performance in the remaining two cases (Fig. 2f and Sup. Table S8). This analysis showed that the automated technique to set the number of hidden units was superior to the manual value selection for this important parameter. The results also showed the capability of DUSC to be a fully automated clustering approach, which can be applied to small datasets (E1, 56 cells) as well as large datasets (MM, 4,139 cells).

### Cluster embedding and visualization

To illustrate the capacity of our approach to preserve the local structure of the data, we generated two-dimensional embeddings for the two largest datasets, MC and MM. Specifically, we applied the t-Distributed stochastic neighbor embedding (t-SNE) method to the latent features generated by DAWN, and compared it to the four other feature learning methods. We considered the MC dataset first, because it was a complex dataset with 3,005 cells and 7 cell types (Fig. 3a). When comparing the embeddings obtained from the original data and after applying the five feature learning methods, the embedding produced from the DAWN-generated latent features showed cells clusters that were the most clearly separated and had smooth elliptical boundaries.

**Figure 3.**
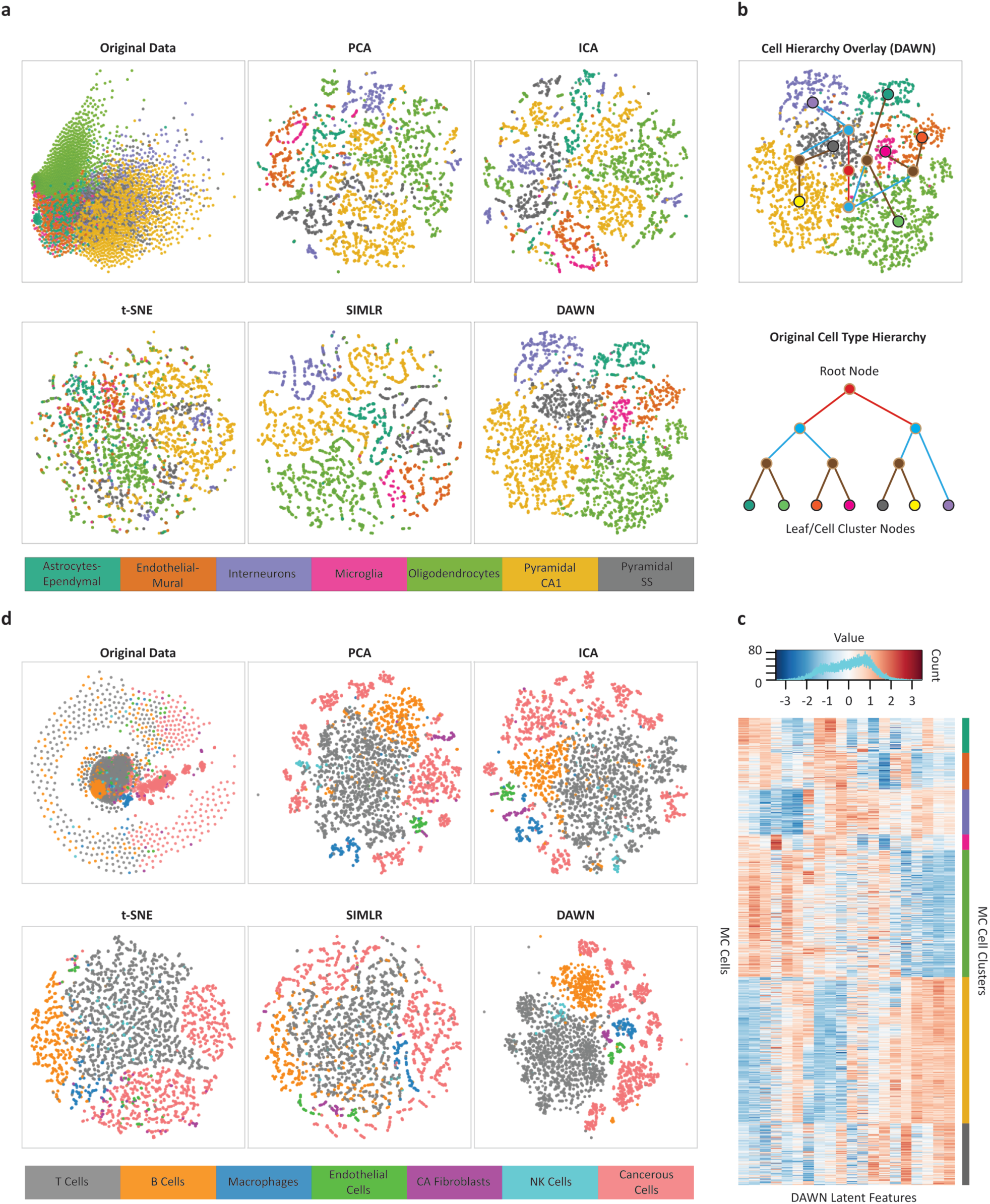
Analysis of clustering performances using visualization approaches. (a) Two-dimensional embedding of the Mouse Cortex (MC) dataset in the original feature space compared with the embeddings of the same dataset in the feature space generated by DAWN and four other feature learning methods, Principle Component Analysis (PCA), Independent Component Analysis (ICA), t-Distributed stochastic neighbor embedding (t-SNE), and Single-cell Interpretation via Multikernel Learning (SIMLR). (b) Hierarchical clustering overlay (top) constructed from the two-dimensional embedding of the DAWN feature space. The hierarchy is created based on the proximities of mass centers of the obtained clusters. The obtained hierarchy is compared to that one of biological cell types (bottom) extracted from the original study [49]. The leaf nodes correspond to the original cell types, while the root and internal nodes correspond to the three other levels obtained during the agglomerative hierarchy. The two-dimensional embedding of the DAWN feature space can recover all but one of defined relationships between the related cell types extracted from the literature. (c) The heatmap of the 20 latent features generated by DAWN on the MC dataset, showing the block structure of the expression profiles of the individual cells grouped by the cell types (bottom). The values of the latent features corresponding to the weights in the hidden layer are distributed in [-3: 3] range (top). (d) Two-dimensional embedding of the Malignant Melanoma (MM) dataset in the original feature space compared with the embeddings of the same dataset in the feature spaces generated by DAWN and four other feature learning methods, PCA, ICA, t-SNE, and SIMLR.

To determine if the biological relationship between the clusters of related cell types could be reflected through the spatial relationship in the 2D embedding, we next created a hierarchical network overlay on the DAWN embedding using the cell type dendrogram obtained from the original study (Fig. 3b). The network topology revealed immediate connections between the clusters corresponding to the more similar cellular types. The more dissimilar clusters were not immediately connected; instead they were connected through the hierarchical nodes and edges in the network, as expected. The obtained network overlay indicated that DAWN preserved the relationships between the cell types during the learning process. The 20 latent features learned by DAWN on the MC dataset were then analyzed using a heatmap representation (Fig. 3c), where the rows represent individual cells, and columns represent the latent features. The heatmap, where the cells were grouped by their types revealed the “block” structural patterns formed by the groups of features, showing that the latent features learned by our method were capable of recovering the intrinsic structure of the original data. The heatmap also shows the orchestrated work of hidden neurons to learn the complementary patterns.

Finally, we obtained the two-dimensional embeddings for the MM dataset (Fig. 3d), another complex dataset with a high variation in the cluster size (52-2,068 cells). We found that DAWN was the only feature learning method capable of producing compact and well-separated clusters, where the two major cell types, *i.e.*, cancerous and non-cancerous cells, were separated with no overlap. The sub-types of non-cancerous cells were also well-separated, with the only exception being natural killer (NK) cells (52 cells), which partially overlapped with the largest cell cluster of T cells (2,068 cells). This overlap could be explained by the disproportionately small size of the NK cluster and the substantial similarity between NK cells and T cells [56].

## Discussion

In this work, we have presented DUSC, a new hybrid approach for accurate clustering of single-cell transcriptomics data. Rapid progress in the development of scRNA-seq technologies urges the advancement of accurate methods for analyzing the single-cell transcriptomics data [57]. One of the first tasks for such analysis is extracting the common patterns shared between cell populations by clustering the cells together based on their expression profiles. The process of clustering, ideally, can help in answering two questions: (1) what is the biological reason that the cells are grouped together (*e.g.*, a shared cellular type), and (2) what are the biological constituents found in the scRNA-seq data that determine the similarity between the cells from the same cluster (*e.g.*, expression values for a set of the overexpressed genes). An important advantage of the clustering methods is their power to extract novel, previously unseen similarity patterns, which leads to the discovery of new cell types [58], spatial cellular compartmentalization in disease and healthy tissues [59], subpopulations of cells from different developmental stages [60], and other cellular states. However, the clustering accuracy, in spite of being continuously tackled by the recent methods, has remained substantially lower when compared to the supervised learning, or classification, methods. Classification methods, in turn, are designed to handle data from the cellular subpopulations whose representatives have been used during the training stage, and therefore cannot identify novel subpopulations. Another question that has not been fully addressed is the robustness of the class definition based on the scRNA-seq data: Does a class defined by a certain supervised classifier depend on other parameters, such as type of experimental protocol, time of the day, developmental stage, or cell location in the tissue?

DUSC improves the clustering accuracy by (i) leveraging a new deep learning architecture, DAWN, which is resilient to the inherent noise in the single-cell data and generates the data representation with automated feature learning, thus efficiently capturing structural patterns of the data, and (ii) pairing this reduced representation with the model-based EM clustering. In particular, DUSC generates more accurate clusters compared to the clustering algorithms alone and is better than four classical and state-of-the-art feature learning methods integrated with the clustering algorithms. Furthermore, our method achieves a comparable performance with a state-of-the-art supervised learning approach. The novel neuronal approximation implemented in the denoising autoencoder simplifies the optimization process for the most important hyper-parameter in the deep architecture— the number of hidden neurons. The simplicity of using DAWN is thus comparable to PCA, and the utility of the newly learnt features is illustrated by the better visualization of large scRNA-seq datasets when using a two-dimensional embedding. Lastly, our multi-tiered assessment reveals the dependence of clustering performance on the dataset complexity, as defined by an information-theoretic metric, which is due to the size balance of the subpopulations in the dataset.

Our next step is to improve the scalability of DUSC for the very large datasets containing 100,000+ cells, which are highly heterogeneous and may include a certain cell type hierarchy [61]. We also plan to evaluate if a deeper architecture can improve the feature learning on the massive datasets. An even more challenging task is to improve feature learning for the highly imbalanced data, *e.g.*, to be able to detect cell subpopulations of disproportionally small sizes, which would either be absorbed by a larger cluster or identified as noise and removed from the analysis by the traditional methods. We have seen this scenario in the MM dataset, the non-cancerous subtypes vary in sample size from 52 cells to 2,068 cells, which affects the performance of all considered methods. Another interesting application of DUSC is to analyze time-sensitive scRNA-seq data of cell differentiation [62]. In summary, we believe that DUSC will provide life scientists and clinical researchers a more accurate tool for single-cell data analysis, ultimately leading to deeper insights in our understanding of the cellular atlas of living organisms, as well as improved patient diagnostics treatment. DUSC is implemented as an open-source tool freely available to researchers through GitHub: https://github.com/KorkinLab/DUSC.

## Acknowledgements

We would like to thank Prof. Eibe Frank for helpful information regarding the functionality of the Weka data mining tool.

## Author Contributions

DK, SS and NTJ conceived the idea. SS and NTJ collected the scRNA-seq datasets. SS implemented the method DUSC. SS, NTJ and DK analyzed the results. SS and DK wrote the manuscript, with all authors contributing to the manuscript discussion and editing.

## Competing Financial Interests

The authors declare no competing financial or non-financial interests.

