## Supplementary Figures and Tables for "A Hybrid Deep Clustering Approach for Robust Cell Type Profiling Using Single-cell RNA-seq Data"

**Supplementary Information**

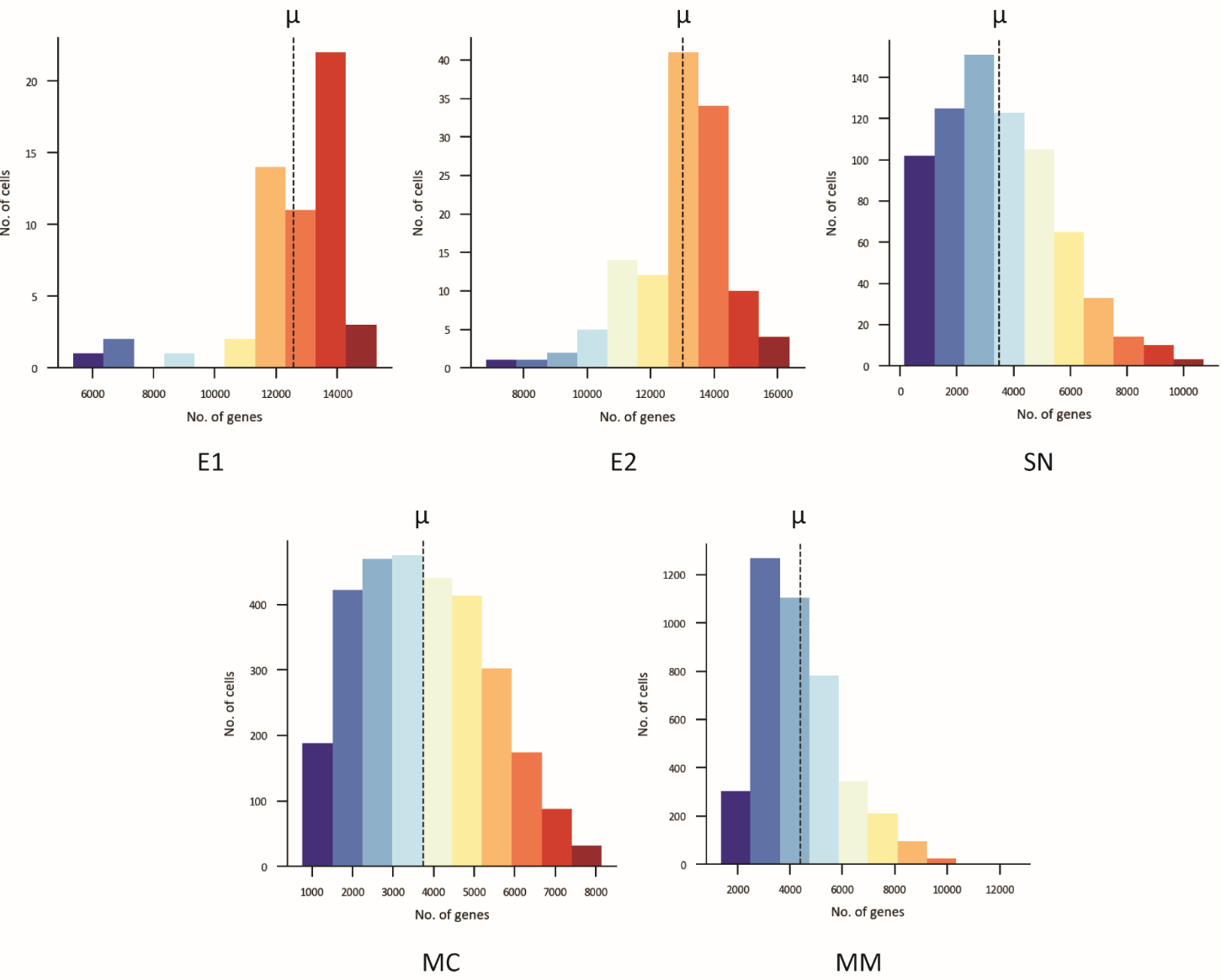

**Supplementary Figure S1:** Number of genes quantified per cell for each scRNA-seq dataset. The two embryonic datasets, E1 and E2 respectively, have fewer samples sequenced compared to the other three datasets, but they have significantly more genes quantified, with an average of 13K genes. The datasets SN, MC and MM on the other hand have an average of about 4K genes quantified.

**Supplementary Table S1:** Number of contiguous zero valued genes/features removed in each scRNA-seq dataset. The original feature set and the resulting feature set after removing zero valued features (ZVF) during the pre-processing phase of DUSC. Note: ERCC genes were removed manually.

| **Dataset** | **Original feature set** | **No. of zero valued features** | **Resulting feature set** |
| --- | --- | --- | --- |
| E1 | 25,737 | 51 | 25,686 |
| E2 | 41,388 | 13,241 | 28,147 |
| SN | 25,334 | 5,710 | 19,624 |
| MC | 19,972 | 0 | 19,972 |
| MM | 23,686 | 904 | 22,782 |

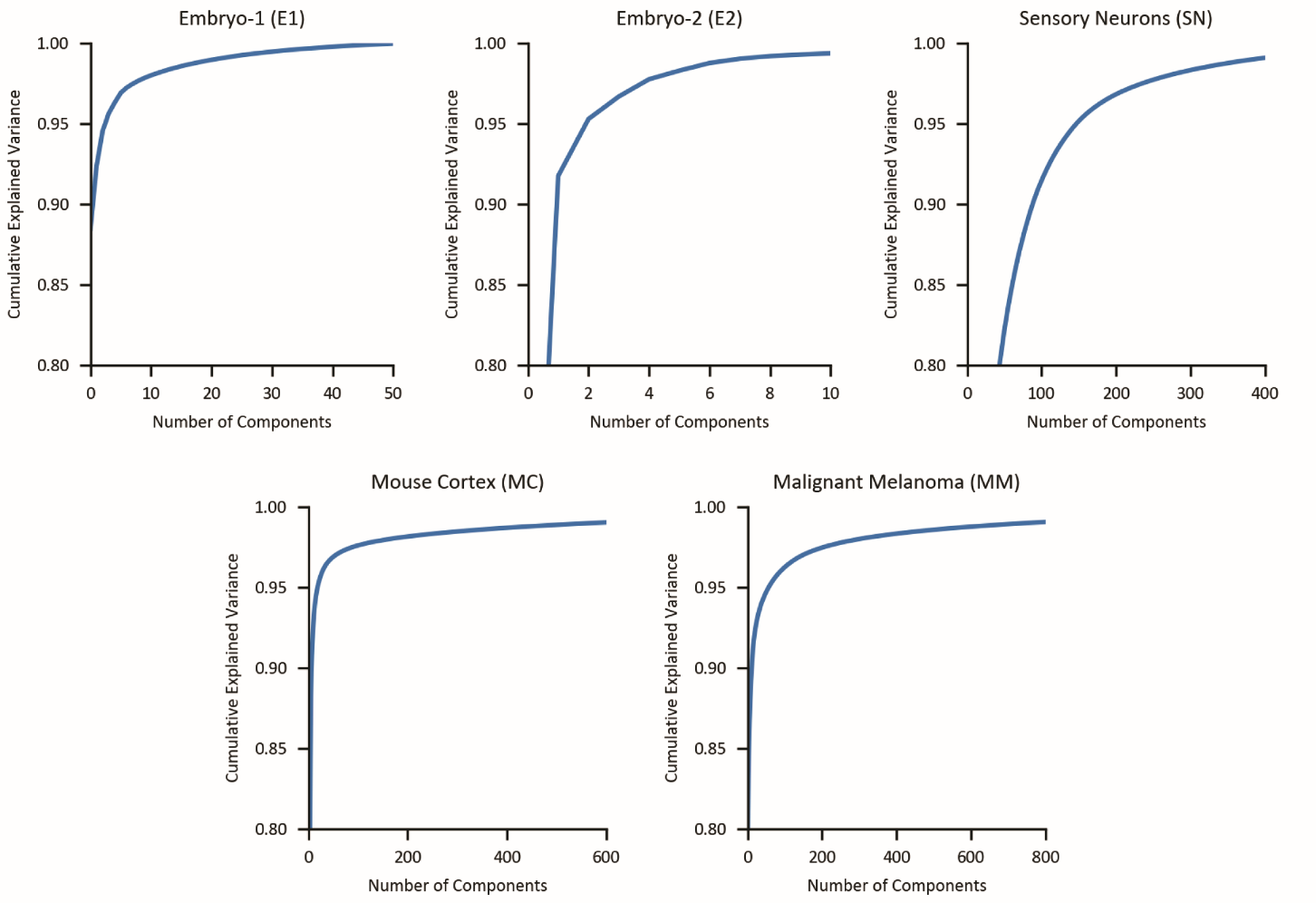

**Supplementary Figure S2:** Principal Component Analysis (PCA) of the scRNA-seq datasets. Since PCA is necessary for ICA and DAWN, detailed plots of variance and components are shown here. The cumulative variance is plotted as a function of the number of principal components that explain the required amount of variance. The Y-axes are initialized to 0.8, since the first component explains at least 0.8 of the variance in each dataset.

**Supplementary Table S2:** Details of clustering and classification accuracies in Fig. 2a. The table shows the accuracies for the proposed method DUSC, the basic clustering algorithms K-means (KM) and Expectation-Maximization (EM), and the supervised learning method Random Forest (RF) respectively.

| **Dataset** | **DUSC** | **KM** | **EM** | **RF** |
| --- | --- | --- | --- | --- |
| E1 | 0.9464 | 0.6607 | 0.9643 | 0.9464 |
| E2 | 0.9597 | 0.6855 | 0.6774 | 0.8790 |
| SN-i | 0.9001 | 0.5321 | 0.6224 | 0.8796 |
| SN-ii | 0.6895 | 0.4523 | 0.5253 | 0.7442 |
| SN-iii | 0.7168 | 0.4596 | 0.4049 | 0.7100 |
| MC | 0.8136 | 0.5398 | 0.4656 | 0.9241 |
| MM-i | 0.9855 | 0.9551 | 0.9650 | 0.9867 |
| MM-ii | 0.6424 | 0.4494 | 0.6202 | 0.9266 |

**Supplementary Figure S3:** Effects of Data Balance on Accuracies for basic clustering methods. This figure bridges the analysis between Fig. 2a, which shows DUSC and the basic clustering methods (KM and EM), and Fig. 2b, which shows the effect of data balance on DUSC. The accuracies and data balance values for DUSC, KM, and EM are shown across different datasets. We see that in datasets where lower balance exists between clusters (*e.g.*, MM-ii, E2, and SN-iii), there is a loss of accuracy by all methods (except for DUSC in E2).

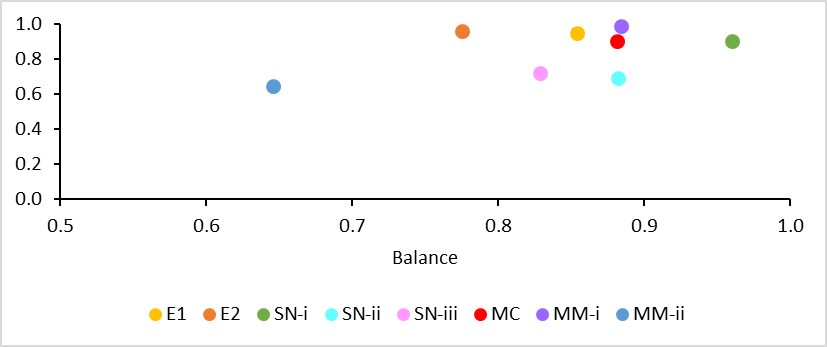

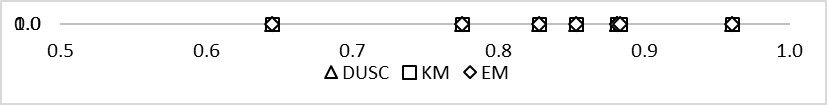

Accuracy

| Data Balance | DUSC | KM | EM |
| --- | --- | --- | --- |
| 0.8541 | 0.9464 | 0.6607 | 0.9643 |
| 0.7756 | 0.9597 | 0.6855 | 0.6774 |
| 0.9607 | 0.9001 | 0.5321 | 0.6224 |
| 0.8821 | 0.6895 | 0.4523 | 0.5253 |
| 0.8289 | 0.7168 | 0.4596 | 0.4049 |
| 0.8819 | 0.8136 | 0.5398 | 0.4656 |
| 0.8843 | 0.9855 | 0.9551 | 0.965 |
| 0.6456 | 0.6424 | 0.4494 | 0.6202 |

**Supplementary Table S3:** Data balance and clustering accuracies shown in Sup. Fig. S3. The data balance for each dataset is listed first and is used in Fig. 2b and Sup. Fig. S3. The clustering accuracies for DUSC, KM, and EM are then noted.

|  | **Balance** | **Accuracy** | | |
| --- | --- | --- | --- | --- |
| **Datasets** |  | **DUSC** | **KM** | **EM** |
| E1 | 0.8541 | 0.9464 | 0.6607 | 0.9643 |
| E2 | 0.7756 | 0.9597 | 0.6855 | 0.6774 |
| SN-i | 0.9607 | 0.9001 | 0.5321 | 0.6224 |
| SN-ii | 0.8821 | 0.6895 | 0.4523 | 0.5253 |
| SN-iii | 0.8289 | 0.7168 | 0.4596 | 0.4049 |
| MC | 0.8819 | 0.8136 | 0.5398 | 0.4656 |
| MM-i | 0.8843 | 0.9855 | 0.9551 | 0.9650 |
| MM-ii | 0.6456 | 0.6424 | 0.4494 | 0.6202 |

**Supplementary Table S4:** Details of clustering accuracies shown in Fig. 2d. The below table includes the accuracies for our feature learning method DAWN, and the other feature learning methods, *i.e.*, SIMLR, PCA, ICA, and t-SNE. In this scenario, all feature learning methods, including DAWN, have been integrated with Expectation-Maximization (EM) clustering. The data allow one to show the impact of EM on the final clusters across all feature learning methods.

| **Dataset** | **DAWN** | **SIMLR** | **PCA** | **ICA** | **t-SNE** |
| --- | --- | --- | --- | --- | --- |
| E1 | 0.9464 | 0.7500 | 0.7679 | 0.8036 | 0.7143 |
| E2 | 0.9597 | 0.5887 | 0.5806 | 0.4516 | 0.5242 |
| SN-i | 0.9001 | 0.7401 | 0.4364 | 0.4596 | 0.7346 |
| SN-ii | 0.6895 | 0.5773 | 0.4665 | 0.3707 | 0.5923 |
| SN-iii | 0.7168 | 0.4815 | 0.3776 | 0.3899 | 0.4692 |
| MC | 0.8136 | 0.6732 | 0.4839 | 0.5817 | 0.3072 |
| MM-i | 0.9855 | 0.8084 | 0.6376 | 0.5001 | 0.9328 |
| MM-ii | 0.6424 | 0.5574 | 0.6163 | 0.5741 | 0.4671 |

**Supplementary Table S5:** Detailed clustering accuracies shown in Fig. 2e. The below table includes the accuracies for our feature learning method DAWN, and the other feature learning methods, *i.e.*, SIMLR, PCA, ICA, and t-SNE. In this scenario all the feature learning methods, including DAWN have been paired with K-means clustering. The data allow one to show the impact of K-means on the final clusters across all feature learning methods.

| **Dataset** | **DAWN** | **SIMLR** | **PCA** | **ICA** | **t-SNE** |
| --- | --- | --- | --- | --- | --- |
| E1 | 0.6786 | 0.7679 | 0.6429 | 0.6250 | 0.6786 |
| E2 | 0.8871 | 0.5887 | 0.4516 | 0.5081 | 0.4839 |
| SN-i | 0.8399 | 0.7442 | 0.3051 | 0.3598 | 0.5718 |
| SN-ii | 0.5034 | 0.5390 | 0.3871 | 0.3338 | 0.4897 |
| SN-iii | 0.4555 | 0.5116 | 0.3242 | 0.3133 | 0.4131 |
| MC | 0.9012 | 0.6353 | 0.5258 | 0.4483 | 0.4775 |
| MM-i | 0.9657 | 0.8094 | 0.9300 | 0.5849 | 0.9570 |
| MM-ii | 0.5980 | 0.6378 | 0.5878 | 0.3965 | 0.4571 |

**Supplementary Table S6:** Number of hidden neurons selected by DAWN for the scRNA-seq dataset. During the feature learning phase, DAWN sets the number of hidden neurons (*i.e.,* the number of features for learning) based on the analysis of each dataset.

| **Dataset** | **N of input features to DAWN** | **Hidden neurons set by DAWN** |
| --- | --- | --- |
| E1 | 25,686 | 4 |
| E2 | 28,147 | 3 |
| SN | 19,624 | 148 |
| MC | 19,972 | 20 |
| MM | 22,782 | 58 |

**Supplementary Table S7:** Specific values for feature compression achieved by DAWN as shown in Fig. 2c. The feature compression values are computed using the values of the column “Hidden neurons set by DAWN” from Sup. Table S6.

| **Data** | **Feature Compression** |
| --- | --- |
| E1 | 0.9998 |
| E2 | 0.9999 |
| SN | 0.9942 |
| MC | 0.9989 |
| MM | 0.9976 |

**Supplementary Table S8:** Clustering accuracies shown in Fig. 2g where different configurations of the DUSC pipeline are shown. The denoising autoencoder with neuronal approximator (DAWN) is compared against two other common neuronal values for the DAE’s hidden layer, *i.e.*, 50 and 100 respectively.

| **Dataset** | **DUSC(DAWN)** | **DUSC(DAE-50)** | **DUSC(DAE-100)** |
| --- | --- | --- | --- |
| E1 | 0.9464 | 0.6964 | 0.7679 |
| E2 | 0.9597 | 0.5726 | 0.5968 |
| SN-i | 0.9001 | 0.6553 | 0.8112 |
| SN-ii | 0.6895 | 0.5595 | 0.5663 |
| SN-iii | 0.7168 | 0.4747 | 0.4145 |
| MC | 0.8136 | 0.7171 | 0.7657 |
| MM-i | 0.9855 | 0.9862 | 0.9652 |
| MM-ii | 0.6424 | 0.6560 | 0.5139 |
